# Growth sensitivity to water availability as potential indicator of drought-induced tree mortality in Mediterranean *Pinus sylvestris* forests

**DOI:** 10.1101/2023.05.18.541207

**Authors:** Asier Herrero, Raquel González-Gascueña, Patricia González-Díaz, Paloma Ruiz-Benito, Enrique Andivia

## Abstract

Drought-associated tree mortality has worldwide increased in the last decades, impacting structure and functioning of forest ecosystems, with implications for energy, carbon and water fluxes. However, the understanding of the factors underlying this mortality are still limited, especially at stand scale. We aim to identify the factors that triggered the mortality of the widely distributed *Pinus sylvestris* in an extensive forest area in central Spain. We compared radial growth patterns in pairs of live and recently dead individuals that co-occur in close proximity and present similar age and size, thereby isolating the effects of size and environment from the mortality process. Temporal dynamics of growth, growth synchrony, and growth sensitivity to water availability (P-PET) were compared between live and recently dead trees. Over the last 50 years, we observed an increase in the growth synchrony and sensitivity to water availability as drought conditions intensified consistent to prior research. However, no differences were found in radial growth between live and dead individuals 15 years before mortality, and dead individuals showed lower growth synchrony and sensitivity to water availability than live ones for much of the period studied. This suggests a decoupling between tree’s growth responses and climatic conditions, which could increase vulnerability to hydraulic failure and/or carbon starvation. Overall, our results point to an important role of growth sensitivity to water availability in tree mortality for *P. sylvestris* at its southern distribution limit.

## Introduction

The widespread increase in tree mortality has become one of the most serious threats to forest ecosystems worldwide over the past few decades (Allen et al., 2010; Millar and Stephenson, 2015; Greenwood et al., 2017). Increased aridity, along with extreme drought and heat events have been identified as the main drivers of tree mortality (Williams et al., 2012; Ruiz-Benito et al., 2013; Hammond et al., 2022). However, both human management legacies and biotic agents can interact with climate stress, further increasing tree vulnerability to mortality (Linares et al., 2010; Anderegg et al., 2015; Sangüesa-Barreda et al., 2015; McDowell et al. 2022). The impact of increasing tree mortality, particularly in the form of massive mortality events, extends beyond individual trees and can have farreaching effects on ecological communities, ecosystem functions and services, and land-climate interactions (Anderegg et al., 2013). High tree mortality can alter species composition and distribution, and ultimately shifts forest communities towards higher dominancy of shrub species (Allen and Breshears, 1998; Peñuelas et al., 2007; Herrero et al., 2013; Herrero and Zamora, 2014; Allen et al., 2015; Ruiz-Benito et al., 2017). The loss of tree cover can also have significant implications for energy, carbon and water fluxes, and nutrient cycling (Hughes et al., 2006; Allen, 2007, Royer et al., 2011; Xiong et al., 2011; Adams et al., 2012, Anderegg et al., 2013).

Despite the current importance of tree mortality in forest functioning and resilience, the understanding of underlying mechanisms and the development of robust early-warning indicators remain limited due to complex interactions between drivers and involved processes (Camarero et al., 2015; Cailleret et al., 2017; Hartman et al., 2018; McDowell et al., 2022). Tree mortality has been widely linked to hydraulic failure and carbon starvation, with evidence suggesting that they are not mutually exclusive but interdependent (McDowell et al., 2011). In this regard, dendroecology provides valuable information to unravel the physiological mechanisms behind tree mortality and to develop early warning indicators (Cailleret et al., 2017; Hartman et al., 2018). Dendroecological studies allow for easy observation on growth rates, widely used as an indicator of tree vitality (Dobbertin, 2005), for almost the entire lifespan of sampled individuals across extensive areas, which is key for assessing climatic effects on tree functioning. Additionally, synchrony in tree radial growth has been widely used as a sign of forest vulnerability to climate change, as high coincidence in growth patterns between individuals may reflect the strong impact of climatic stressors (Boden et al., 2014; Shestakova et al., 2016). Overall, the study of temporal changes in tree growth responses to drought and warming can provide insights into individual tree predisposition to mortality.

The comparison of growth patterns between co-occurring dead and surviving trees has become a common method for modeling and predicting tree mortality (Pedersen, 1998; Ogle et al., 2000; Cailleret et al., 2017). In many cases, a decline in growth prior to mortality has been observed, although the magnitude and duration of this decline vary depending on species’ functional characteristics and factors driving mortality (Gea-Izquierdo et al., 2014, 2019; Cailleret et al., 2017). Thus, growth patterns prior to mortality can provide valuable information on the physiological mechanisms underlying the mortality process, as changes in tree function (e.g. hydraulic conductivity) and structure (e.g. leaf area) affecting tree growth often precede mortality (McDowell et al., 2011; Seidl et al., 2011). Growth synchrony can also serve as an early warning indicator of mortality. For example, Cailleret et al. (2019) showed a decrease in the synchrony of gymnosperm species 20 years before mortality (Cailleret et al., 2019). However, to make live/dead comparisons more robust, it is essential to standardize variables that may affect radial growth and its variability, such as tree size and biotic and abiotic modulators (e.g. Muñoz-Galvez et al., 2021). Studies that compare growth patterns between live and dead trees while controlling for these key variables may provide new insights into possible growth-based early warning indicators. Given the wide variety of species and population responses, types of climatic stress, and combinations of mortality drivers (Cailleret et al., 2017), such studies have the potential to add important dimensions to our existing knowledge.

Tree species at the limits of their distribution often face particularly adverse ecological conditions, such as temperate or boreal species in drought-prone Mediterranean areas (Hampe and Petit, 2005; Hampe and Jump, 2011). Climate change scenarios project a worrying increase in the magnitude and frequency of drought events in Mediterranean areas (IPCC, 2022), making these species especially vulnerable to drought-induced mortality (Galiano et al., 2010, Herrero et al., 2013; Vila-Cabrera et al., 2013). Furthermore, reforestation and afforestation, and the abandonment of traditional forest management have increased the density of many Mediterranean forests, leading to increased inter-individual competition for water resources (Linares et al., 2010; Vila-Cabrera et al., 2013) and ultimately to forest decline and mortality events (Martínez-Vilalta and Piñol, 2002; Galiano et al., 2010; Vila-Cabrera et al., 2013; Gazol and Camarero, 2022). In this context, there is a critical need to study growth-based early-warning indicators of mortality in tree species’ rear-edge populations in Mediterranean areas, which can aid in implementing forest management and biodiversity conservation actions for the adaptation of these ecosystems to climate change.

In this study, we aimed to investigate the potential for growth-based mortality indicators in Scots pine (*Pinus sylvestris* L.) trees in a high-density forest in Central Spain. *P. sylvestris* is an economically important and widely distributed boreo-alpine species in Europe, with its southernmost limit in the Iberian Peninsula where it has experienced recent mortality events (Galiano et al., 2010; Vila-Cabrera et al., 2013). For this, we (i) evaluated the effects of intrinsic (size), biotic (competition), and abiotic (water availability) factors on the growth rates of both dead and surviving trees, (ii) analyzed the temporal pattern of growth sensitivity to water availability for dead and surviving trees, and (iii) quantified growth synchrony. By doing so, we sought to gain insights into the mechanisms underlying tree mortality and help to develop a reliable growth-based mortality indicator for *P. sylvestris* in Mediterranean areas. In accordance with previous findings in gymnosperm species, we expect lower growth rates in dead than in surviving trees before mortality, likely due to their drought response-related functional characteristics (Cailleret et al., 2017). Furthermore, we anticipate that competition will have a greater impact on growth in dead trees, which could intensify the adverse impact of drought (Linares et al., 2010). Finally, we also expect a decrease in growth synchrony in dead trees before mortality (Cailleret et al., 2019), suggesting a mismatch between growth and high-frequency climatic fluctuations.

## Material and methods

### Study area

The study area is an extensive pine forest located in the Alto Tajo Natural Park of central Spain at an elevation range of 1450-1600 m a.s.l. (40° 39’ 6.8”N, 1° 41’ 53.1”W). This forest experienced a widespread tree mortality event during 2016 and 2017. No sign of biotic agents was detected. The forest area is dominated by *P. sylvestris* and *P. nigra* Arnold, with *P. pinaster* Ait in some areas. The soil type in the area is primarily calcium cambisols (Martín-Moreno et al., 2014). The study area has a continental Mediterranean climate with cold winters and mild and dry summers, with a mean annual temperature of 10.5 °C and a mean annual total precipitation of 468 mm (period 1948-2017), according to data from a meteorological station of AEMET (Spanish Meteorological Agency) located at Molina de Aragón, which is situated 25 km from the study area.

### Sampling design and dendroecological methods

In October 2017, we sampled 39 pairs of trees within the study area. For each pair, one tree was observed to be in a healthy state, while the other was dead, but with dry needles still present in the crown indicating recent mortality. The sampled pairs had similar size and they were located no more than 5 meters apart from each other (with an average distance of 3 meters). Two cores were taken from each tree in opposite directions using a Pressler increment borer. The diameter at breast height (DBH) of each sampled tree was also recorded, along with the DBH of every neighboring tree within a circular plot of 10 meters radius measured from the sampled trees.

Cores were processed following standard ecological methods (Fritts, 1976). They were air-dried for 48-72 hours, mounted on wooden supports, and sanded with progressively finer grades of sandpaper to highlight ring-width patterns. All cores were visually cross-dated following the procedures described by Yamaguchi (1991). Cores were scanned at 1,200 dpi resolution (EPSON Perfection v800) and treering width was measured using Image J software (Schneider, 2012). Trees can die during the growing season before ring formation is complete, which induces an incomplete outermost ring. As tree could have died at different timing during the 2017 growing season, we did not consider the last ring of sampled trees (Cailleret et al., 2017). Ring width was converted to basal area increment (BAI, cm^2^ year^-1^) assuming a circular stem cross-section and using the following formula:

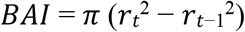

where r_t_ and r_t-1_ are the radius of the tree at the end and at the beginning of a given annual ring, respectively. For each tree and year, the BAI was calculated as an average of the two extracted cores. BAI is a meaningful indicator of tree growth as it removes the variation in growth due to increasing circumference, making it more related to biomass increment than ring width (Biondi and Qeadan, 2008).

To quantify inter-individual competition in sampled trees, we calculated the Lorimer’s competition index (LCI, Lorimer, 1983) using the following formula:

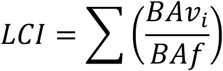

where *BA*v_i_ is the basal area of each neighbor tree within the circular plot of 10 m radius and *BA*f is the basal area of the sampled tree.

### Extreme droughts

To quantify the annual water availability, we calculated the annual water balance by subtracting the annual potential evapotranspiration (PET) from the total annual precipitation (P). The PET was determined following the method proposed by Thornthwaite (1948). To account for the influence of previous year conditions on the current growing season, we calculated annual water balance from October of the previous year to September of the present year (Madrigal-Gonzalez et al., 2018). We identified the main extreme droughts during the study period as those years with annual P-PET below the first percentile of the P-PET series (Steckel et al., 2020).

### Data analysis

We used a linear mixed-effect model to compare BAI series between live and dead trees and assess the effect of intrinsic, biotic and abiotic factors on growth. We used data from the 1948-2017 period to increase sample size and statistical power of the analysis. We included tree identity nested within tree-pairs as a random effect to account for the repeated measurements on the same individuals and for data on paired individuals. We also used an autoregressive structure to consider the temporal autocorrelation of successive annual growth increments (Pinheiro et al., 2018). Size (DBH of the current year), state (live/dead), competition (LCI), water balance (P-PET) and the interactions state × water balance and state × competition were considered as fixed effects in the model. Size was included as a second-degree polynomial to allow for nonlinear responses of growth to variations in size (Sumida et al., 2013). Finally, to test for temporal changes in the effect of environmental factors (particularly water balance) and size on tree growth, the model was fitted to 5-year lagged windows of 20 years (Díaz-Martínez et al., 2023; i.e., 11 time-windows of 20 years with a starting year from 1948 to 1998).

To assess growth synchrony within dead and surviving trees we followed the procedure described by Alday et al. (2018). Briefly, we used variance-covariance mixed modelling using year as random term and calculated the mean inter-chronology correlation (i.e. growth synchrony) as the common interannual growth variance to all dead and surviving trees. To do this, we tested different variance-covariance structures (see Alday et al., 2018 for a complete overview of variance-covariance structures) that allow to characterize synchrony for each level of the grouping variable (i.e. dead and surviving trees). The best variance-covariance structure was selected using the Akaike Information Criterion (AIC, Zuur et al., 2009). The temporal changes in growth synchrony for dead and surviving trees were also assessed using 20-year intervals with a 5-year moving window.

All statistical analyses were done in R 3.5.2 (R Core Team, 2018) using the *nlme* (Pinheiro et al., 2018), *MuMn* (Barton, 2019) and *R DendroSync* (Alday et al., 2018) packages.

## Results

### Characteristics of sampled trees

Considering raw growth data (BAI) for the 1900-2017 period, live and dead trees showed similar patters along the 1900-2017 period, yet live trees showed lower mean BAI than dead trees (Table 1), especially between 1930 and 1980 (Fig. 1). After 80s, live and dead trees showed similar growth rates, except in the last years when dead trees also showed greater growth than live trees (Fig. 1). Age, size (DBH) and competition index (LCI) were similar for both live and dead trees (Table 1). Sampled trees presented an average age of 130 years and average DBH of 34 cm (Table 1).

**Table 1.** Characteristics of the sampled trees (mean ± SD). BAI: Basal Area Increment (1900-2017); DBH: Diameter at Breast Height (1.3m); BA: Basal Area of the 10 m radius neighbourhood around the sampled tree; and LCI: Lorimer’s Competition Index.

|  | All | Live | Dead |
| --- | --- | --- | --- |
| **BAI (cm<sup>2</sup>)** | 5.33 ± 4.42 | 4.81 ± 3.89 | 5.84 ± 4.84 |
| **Age at breast height (years)** | 129.96 ± 40.53 | 127 ± 44 | 133 ± 37 |
| **DBH (cm)** | 33.97 ± 7.17 | 33.51 ± 7.3 | 34.30 ± 7.14 |
| **BA (m<sup>2</sup> ha<sup>-1</sup>)** | 26.93 ± 10.88 | 26.83 ± 11.64 | 27.27 ± 8.03 |
| **LCI** | 13.79 ± 10.33 | 14.15 ± 11.24 | 12.50 ± 6.34 |

**Figure 1.**
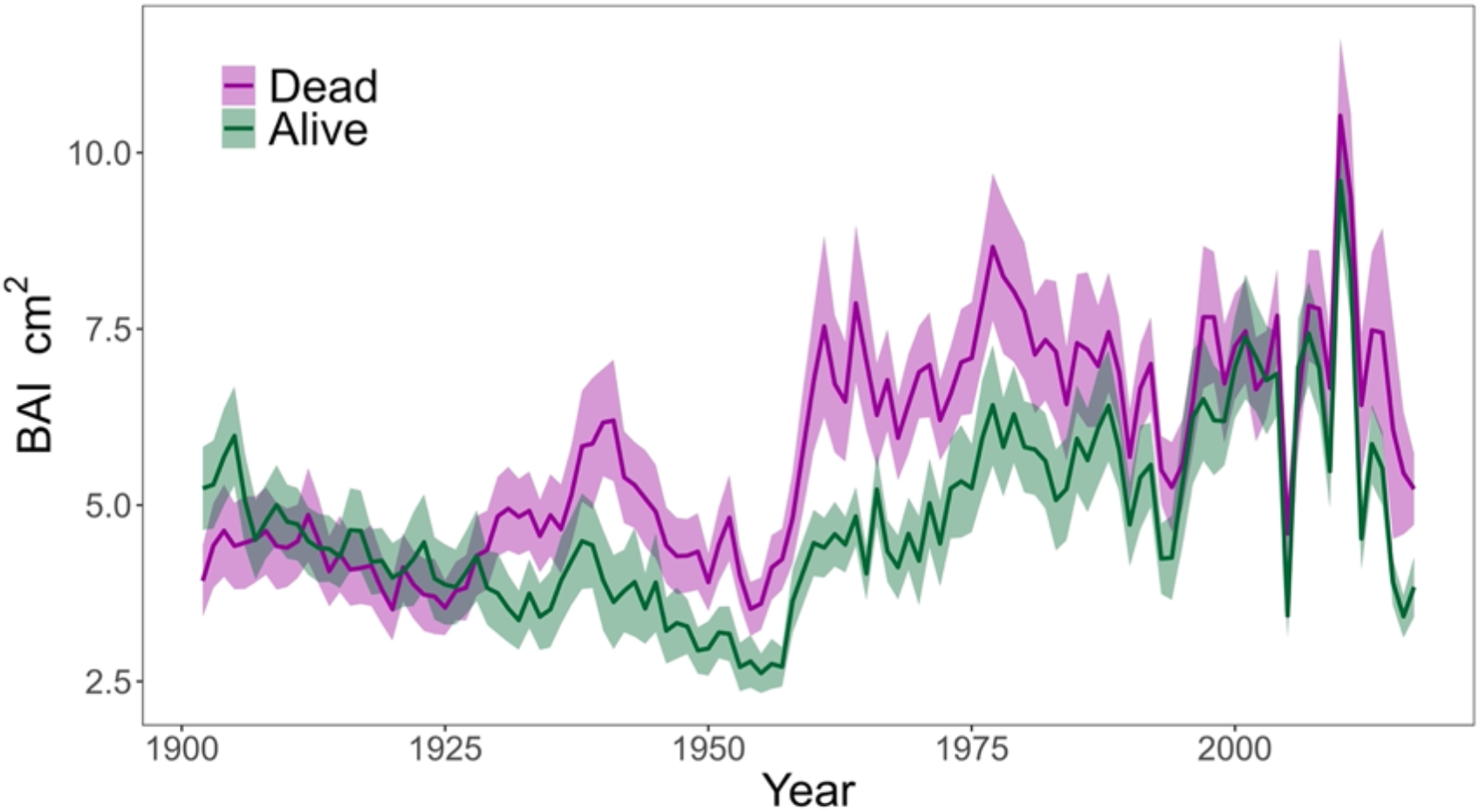
Basal Area Increment (BAI, cm^2^, ± SE) of live and dead trees evaluated for the 1900-2017 period.

### Effect of size, competition and climate on growth

Tree radial growth was significantly influenced by tree size, competition index and the interaction state × water balance. No significant differences were found on tree growth between live and dead trees for the 1948-2017 period (Table 2).

**Table 2.** Results (F and p) from the growth model fitted to evaluate the effect of the following factors: tree size as a second-degree polynomial (Poly (size)), water balance (P-PET), competition index (LCI) and interactions state × water balance and state × competition.

|  | F | P |
| --- | --- | --- |
| **Poly (size)** | 86.7452 | < 0.0001 |
| **State** | 1.9817 | 0.1681 |
| **P-PET** | 207.1174 | < 0.0001 |
| **LCI** | 7.3575 | 0.0103 |
| **State\*P-PET** | 8.9729 | 0.0028 |
| **State\*LCI** | 0.2440 | 0.6244 |

The competition index had a negative effect on growth, whereas the effect of the water balance was positive (Table S1). Dead trees showed significantly less sensitivity to water balance (lower slope) than live trees. Further, when the water availability was low, dead trees showed greater growth than live trees (Fig. 2).

**Figure 2.**
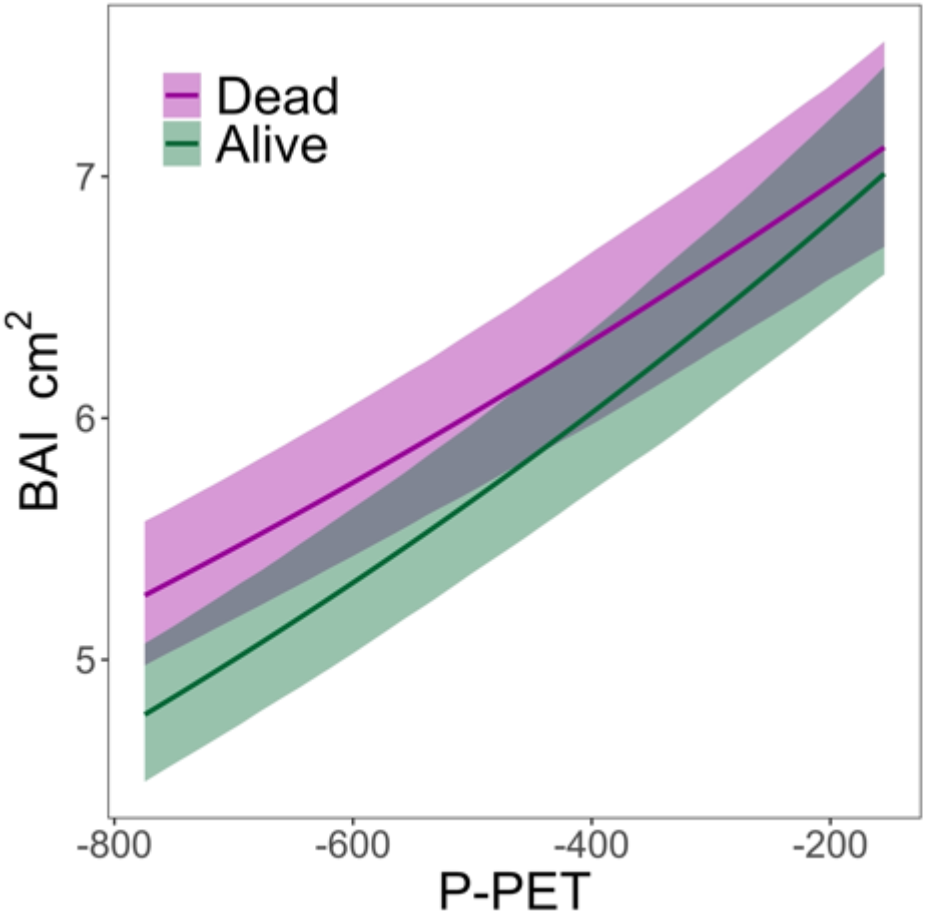
Growth response (BAI, cm^2^, ± SE) to water balance (P-PET, mm, ± SE) for the period 1948-2017 in dead and live trees.

### Temporal changes in growth patterns

Time windows analysis showed similar growth patterns between live and dead trees for the studied period (Fig. 3).

**Figure 3.**
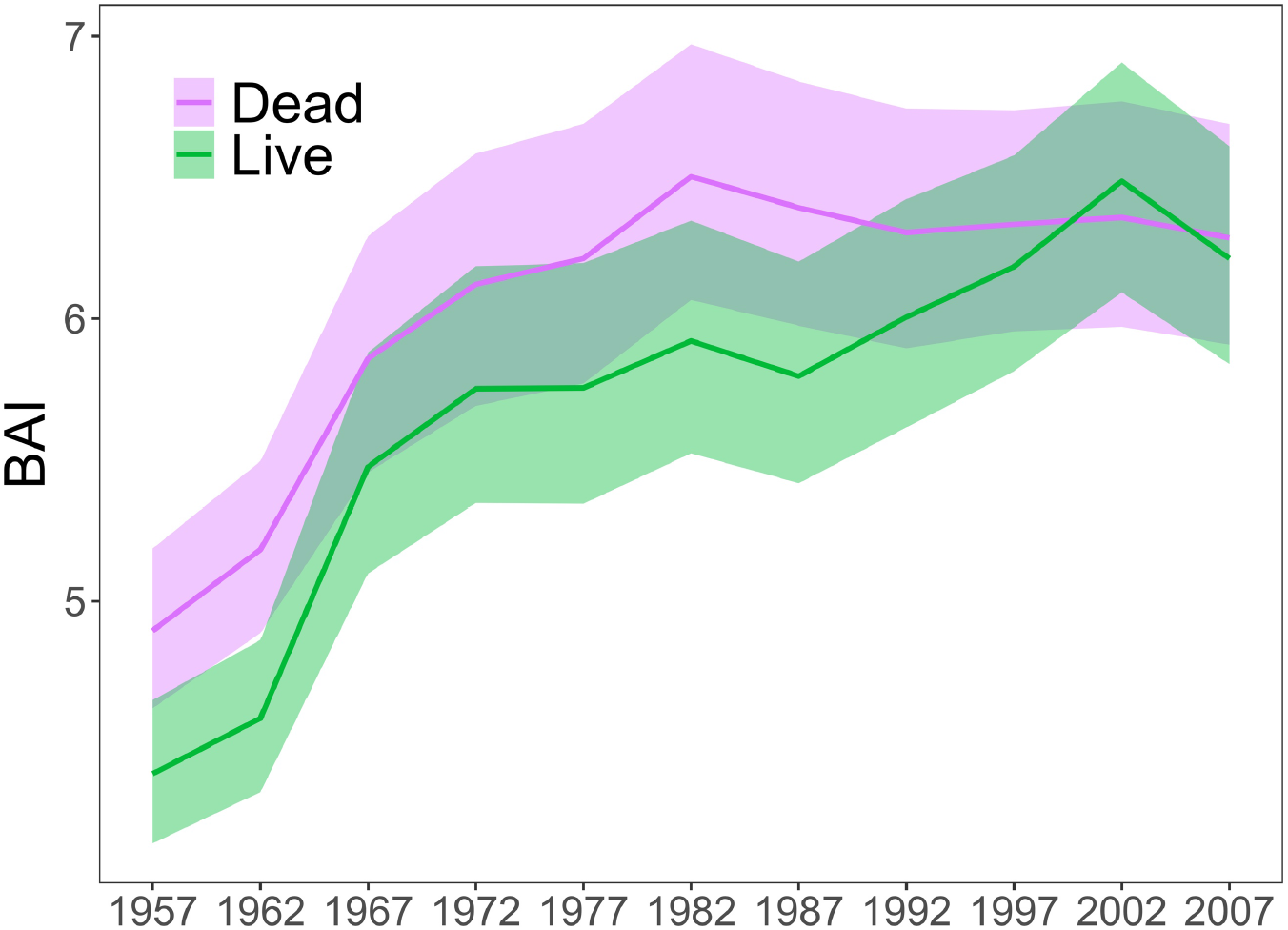
Temporal growth evolution (BAI, cm^2^, ± SE) in live and dead trees for the 1947-2017 period applying the model at 20-year intervals with a 5-year moving window. The years appearing on the y-axis correspond to the midpoints of each 20-year interval.

Growth sensitivity to water balance increased along the study period, coinciding with a decrease in water availability (Fig. 4). Live trees showed greater sensitivity than dead trees during the first half of the study period. However, these differences disappeared as water balance decreased (Fig. 4). Moreover, dead trees showed lower growth sensitivity to water balance than live ones in the last two 20-year intervals considered.

**Figure 4.**
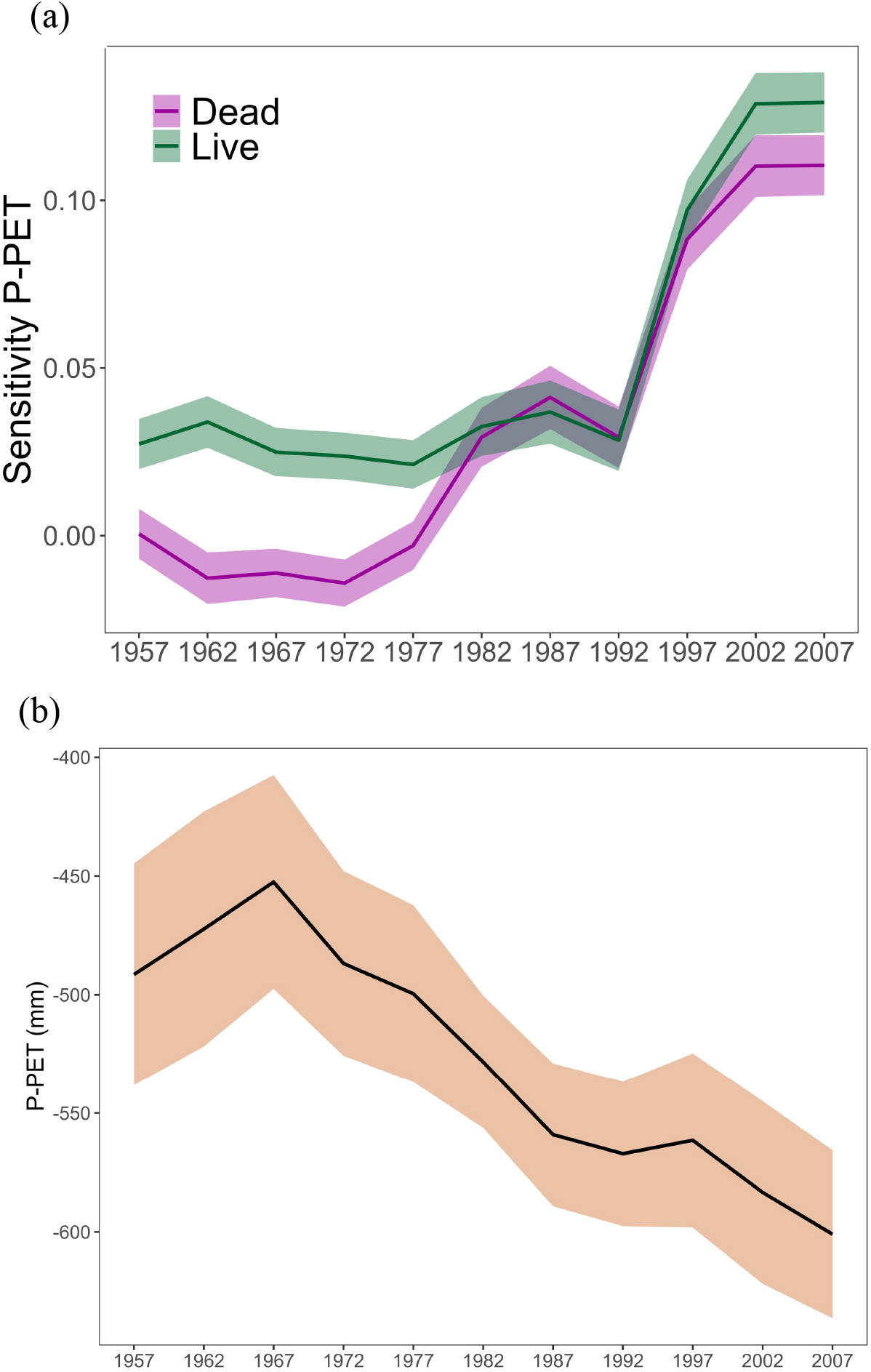
Temporal evolution of (a) growth sensitivity to water balance (i.e., slope of water balance in the growth model) in live and dead trees, and of (b) the water balance (P-PET) for the 1947-2017 period. Shaded areas represent the ± 95% confidence intervals. Data are showed for 20-year intervals with a 5-years moving window. The years appearing on the y-axis correspond to the midpoints of each 20-year interval.

### Growth synchrony

The best-fit variance-covariance structure for calculating synchrony was a heterocedastic matrix with a composite symmetry structure (mHeCS; Table S2). Firstly, the selection of a heterocedastic matrix implies that the magnitude of the common growth signal (i.e. growth synchrony) varied between live and dead trees. Secondly, a matrix with a symmetrical structure suggests the existence of fluctuations in growth that were common for both live and dead trees. Overall, live trees exhibit significantly higher synchrony than dead ones (0.23 ± 0.03 and 0.16 ± 0.02, respectively).

Regarding temporal changes in synchrony, growth synchrony values were relatively low until the last 30 years, when a steep increase in synchrony was recorded (Fig. 5). The synchrony values of live and dead trees diverged markedly during two periods: in the second quarter of the study period and in the last interval analyzed (just before the mortality event) (Fig. 5).

**Figure 5.**
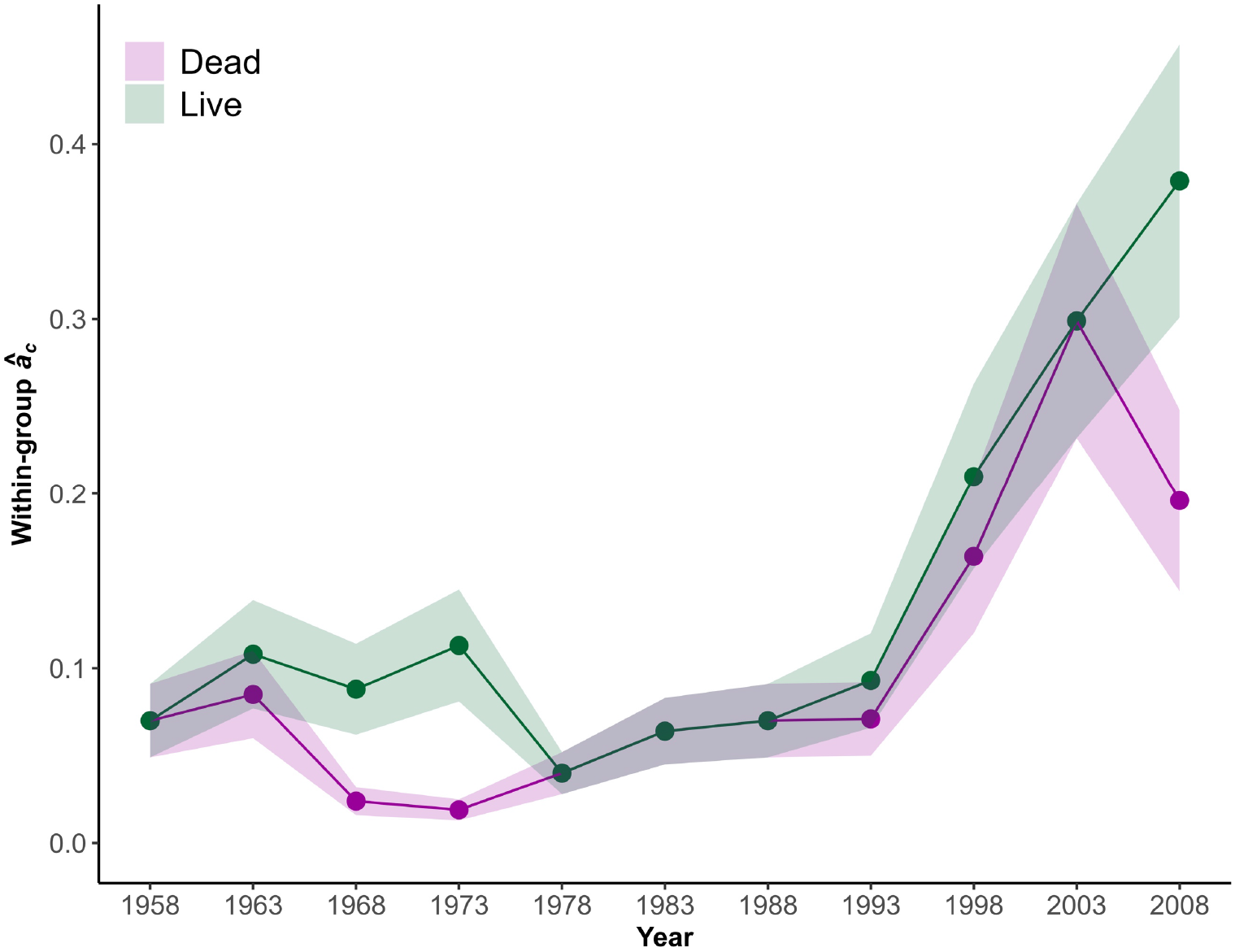
Temporal evolution of growth synchrony for live and dead trees for the 1947-2017 study period. Data are showed for 20-year intervals with a 5-years moving window. The years appearing on the y-axis correspond to the midpoints of each 20-year interval.

## Discussion

Our study revealed significant differences between live and dead trees in growth-related parameters under an increasing aridity context. Both, growth sensitivity to water balance and synchrony were lower in dead than in live trees before mortality. This suggests a decoupling of tree growth from climatic fluctuations in dead trees that may have increased their risk of mortality due to hydraulic failure and/or carbon starvation. Thus, growth sensitivity to water balance should also be considered as an early-warning indicator of mortality in Mediterranean *P. sylvestris* populations located at the southern distribution limit of the species.

Contrary to our first hypothesis, our study did not find lower growth in dead than in live trees prior to mortality, nor a sustained decrease in growth in dead trees. These results are in contrast to those reported by Cailleret et al. (2017) for several gymnosperm species. Nevertheless, our results are consistent with other studies that have found no differences between dead and live trees at stand and regional scales (Ferrenberg et al., 2014; Rowland et al., 2015; Berdanier and Clark, 2016; Herguido et al., 2016). When considering our design based on dead/live tree pairs, there was no significant difference in growth between dead and live trees during the first 50 years of life, ruling out the possibility that mortality was due to a trade-off between early-growth and longevity (Bigler et al., 2007; Bigler and Veblen, 2009). Additionally, the size of live and dead trees was similar. Therefore, we argue that dead trees could have prioritized radial growth until death regardless of environmental stress, or that mortality occurred due to an extreme environmental stress leading to tree death without a previous impact on its carbon balance (Cailleret et al., 2017).

Interestingly, we found that the effect of water balance on growth varied between live and dead trees, with the latter showing higher growth during extremely dry conditions. Thus, under severe drought conditions, dead trees may exhibit a less conservative water use than live trees, which could expose them to a higher risk of hydraulic failure due to xylem embolism (McDowell et al., 2022). Our results contrast with previous studies on *P. sylvestris*, which showed greater sensitivity to drought and hot conditions for dead trees and a sustained decline in growth (Heres et al., 2012). However, our results are consistent with studies on saplings, where *P. sylvetstris* appears unable to down-regulate its height growth under extreme drought conditions (Herrero et al., 2013). These differences in growth responses to drought conditions may be due to intraspecific variability and the intensity and recurrence of drought events in the study area (Cailleret et al., 2017).

Regarding the effect of competition, we did not find support for a greater impact of competition on growth in dead trees. Our results showed that negative effects of competition on tree growth were similar for live and dead trees, which suggests a limited effect of competition on the mortality event. Nonetheless, the consistent negative effect of competition on radial growth of live and dead trees suggests the key role of competition on exacerbating drought impacts on Mediterranean forests (Linares et al., 2010; Vila-Cabrera et al., 2013). However, it should be noted that we used a static competition index measured at sampling, so it can not reflect the effect of competition on the entire growth series. This represents an important limitation in clarifying the role that competition may play in drought-induced growth reductions (Marqués et al., 2021).

The study area experienced a progressive decrease in water balance over time, resulting in reduced water availability for the trees. As drought conditions intensified, sampled trees increased its growth sensitivity to water balance. However, at the beginning and at the end of the study period, dead trees showed lower sensitivity to water balance, indicating potential differences in physiological responses to water stress between live and dead individuals. The gradual increase in aridity may have pushed individuals closer to threshold values where xylem embolism processes occur, which can potentially lead to mortality. Additionally, rising temperatures may increase respiration costs, elevating the risk of carbon starvation during dry periods of low carbon assimilation (Adams et al., 2009; McDowell et al., 2011). Feedbacks between plant hydraulics and carbon balance can also result in synergistic negative effects on tree vigor and health by increasing temperatures and aridity (McDowell et al., 2011). Therefore, in this context of increasing aridity, reduced sensitivity of growth to water balance may increase the risk of drought mortality and could be used as an early-warning signal of tree mortality.

The patterns in growth synchrony observed in our study are similar to those in sensitivity to water balance. We observed an increase in synchrony between individuals as aridity increased, which is consistent with previous studies at local and regional scales (Boden et al., 2014; Shestakova et al., 2016). In fact, increased synchrony is considered an indicator of tree vulnerability to climate change, as it reduces individual variability and resilience capacity at population level (Clark et al., 2012; Tejedor et al., 2020). In line with our third hypothesis, lower synchrony was observed in dead trees both prior to the mortality event and during the first half of the study period. This decrease in synchrony before mortality agrees with previous studies in gymnosperms (Cailleret et al., 2019) and is consistent with the reduction in growth sensitivity to water balance. Both aspects indicate a decoupling between growth and climatic fluctuations, which could be linked to the aforementioned less conservative water use in dead than in live trees (Herrero et al., 2013). Thus, under extreme dry conditions, a higher water expense through a low stomatal responsiveness could increase its vulnerability to cavitation and the risk of mortality from hydraulic failure (McDowell et al., 2008). Thus, growth synchrony should be also considered as a potential indicator for mortality events in seasonally limited areas within *P. sylvestris* distribution.

Overall, our study, which takes advantage of a paired design that minimizes the effects of size and immediate tree environment on potential causes of mortality, places on the table sensitivity to water balance and growth synchrony as potential indicators of drought mortality in southern populations of the widely distributed *P. sylvestris*. To the best of our knowledge, mortality events had not been yet associated with decreases in radial growth sensitivity to water availability. Thus, we recommend considering changes in growth sensitivity to water availability in further studies aimed at elucidating potential mechanisms of tree mortality at local and global scale. Although mortality in individuals that do not show decreases with drought is usually associated with hydraulic failure (Cailleret et al., 2017), carbon starvation cannot be ruled out due to the interdependence of these two processes and the context of increasing temperatures and aridity in Mediterranean and semi-arid ecosystems (IPCC, 2021). Moreover, the quantification of other variables, in addition to radial growth, related to tree vigor (e.g., branch and foliar biomass) and physiological responses to drought (e.g., water use efficiency) as well as a more robust evaluation of competition along tree lifespan may also provide valuable insights into mortality mechanisms and inform forest management for the adaptation to climate change.

## Data availability statement

The data are freely available upon request to the authors.

## Author Contributions

Sampling design and idea: EA, PRB and AH. Field work: EA, PGD, PRB. Sample processing: RGG. Analyses: EA and AH. Manuscript writing: AH. All the authors contributed to the article and approved the submitted version.

## Acknowledgements

We thank Daniel Gómez and Pablo Quiles for field assistance.

## Supplementary Material

**Table S1.** Growth model parameters. It includes the effect of the size as a second-degree polynomial (Poly (size)1 and Poly (size)2), the water balance (P-PET), the competition index (LCI) and the interaction between state and P-PET. The intercept refers to dead trees.

|  | **Value** | **Standard error** | **df** | **t-value** | **p-value** |
| --- | --- | --- | --- | --- | --- |
| **Intercept (state=dead)** | 1.749 | 0.049 | 5152 | 35.320 | 0.0000 |
| **Poly(size)1** | 19.687 | 1.886 | 5152 | 10.434 | 0.0000 |
| **Poly(size)2** | -11.158 | 1.524 | 5152 | -7.319 | 0.0000 |
| **State(live)** | -0.066 | 0.053 | 35 | -1.245 | 0.2213 |
| **P-PET** | 0.037 | 0.005 | 5152 | 8.005 | 0.0000 |
| **LCI** | -0.126 | 0.049 | 35 | -2.533 | 0.0160 |
| **State (live)\*P-PET** | 0.020 | 0.007 | 5152 | 2.995 | 0.0028 |
| **State (live)\*LCI** | 0.026 | 0.054 | 35 | 0.494 | 0.6244 |

**Table S2.** Variance-covariance model comparison for calculating growth synchrony of BAI chronologies of sampled dead and live trees. n: number of observations used in the model fit; df: degrees of freedom related with the number of parameters in the fitted model; AIC: Akaike’s Information Criterion (AIC); AICc: corrected AIC; BIC: Bayesian Information Criterion. The selected model is shown in bold.

| **Model\*** | **n** | **df** | **AIC** | **AICc** | **BIC** |
| --- | --- | --- | --- | --- | --- |
| *Homoscedastic* |  |  |  |  |  |
| mBE | 5372 | 80 | 3300.3 | 3302.7 | 3826.2 |
| mNE | 5372 | 81 | 3383.8 | 3386.3 | 3916.3 |
| mCS | 5372 | 82 | 3303.0 | 3305.6 | 3842.1 |
| mUN | 5372 | 82 | 3293.7 | 3296.3 | 3832.8 |
| *Heteroscedastic* |  |  |  |  |  |
| mBE | 5372 | 80 | 3300.3 | 3302.7 | 3826.2 |
| mHeNE | 5372 | 82 | 3278.9 | 3281.4 | 3818.0 |
| mHeCS | 5372 | 82 | 3198.5 | 3201.0 | 3737.6 |
| mHeUN | 5372 | 83 | 3188.7 | 3191.3 | 3734.4 |

\*Model abbreviations: mBE: Broad Evaluation model; mNE: Narrow Evaluation model; mCS: Compound Symmetry model; mUN: Unstructured model; mHeNE: heteroscedatic variant of mNE; mHeCS: heteroscedatic variant of mCS; mHeUN: heteroscedatic variant of mUN.

## References

Adams, H. D., Guardiola-Claramonte, M., Barron-Gafford, G. A., Villegas, J. C., Breshears, D. D., Zou, C. B., et al. (2009). Temperature sensitivity of drought-induced tree mortality portends increased regional die-off under global-change-type drought. Proc. Natl. Acad. Sci. 106, 7063– 7066. doi: 10.1073/pnas.0901438106.

Alday, J. G., Shestakova, T. A., Resco de Dios, V., and Voltas, J. (2018). DendroSync: An R package to unravel synchrony patterns in tree-ring networks. Dendrochronologia 47, 17–22. doi: https://doi.org/10.1016/j.dendro.2017.12.003.

Allen, C. D., and Breshears, D. D. (1998). Drought-induced shift of a forest-woodland ecotone: Rapid landscape response to climate variation. Proc. Natl. Acad. Sci. U. S. A. 95, 14839–14842.

Allen, C. D. (2007). Interactions across spatial scales among forest dieback, fire, and erosion in northern New Mexico landscapes. Ecosystems 10, 797–808.

Allen, C. D., Macalady, A. K., Chenchouni, H., Bachelet, D., McDowell, N., Vennetier, M., et al. (2010). A global overview of drought and heat-induced tree mortality reveals emerging climate change risks for forests. For. Ecol. Manage. 259, 660–684. https://doi.org/10.1016/j.foreco.2009.09.001.

Allen, C. D., Breshears, D. D., and McDowell, N. G. (2015). On underestimation of global vulnerability to tree mortality and forest die-off from hotter drought in the Anthropocene. Ecosphere 6, art129. doi: 10.1890/es15-00203.1.

Anderegg, W. R. L., Kane, J. M., and Anderegg, L. D. L. (2013). Consequences of widespread tree mortality triggered by drought and temperature stress. Nat. Clim. Chang. 3, 30–36. doi: 10.1038/nclimate1635.

Anderegg, W. R. L., Hicke, J. A., Fisher, R. A., Allen, C. D., Aukema, J., Bentz, B., et al. (2015). Tree mortality from drought, insects, and their interactions in a changing climate. New Phytol. 208, 674–683. doi: https://doi.org/10.1111/nph.13477.

Barton, K. (2009) Mu-MIn: Multi-model inference. R Package Version 0.12.2/r18. http://R-Forge.R-project.org/projects/mumin/

Berdanier, A. B., and Clark, J. S. (2016). Multiyear drought-induced morbidity preceding tree death in southeastern U.S. forests. Ecol. Appl. 26, 17–23. doi: https://doi.org/10.1890/15-0274.

Bigler, C., Gavin, D. G., Gunning, C., and Veblen, T. T. (2007). Drought induces lagged tree mortality in a subalpine forest in the Rocky Mountains. Oikos 116, 1983–1994. doi: https://doi.org/10.1111/j.2007.0030-1299.16034.x.

Bigler, C., and Veblen, T. T. (2009). Increased early growth rates decrease longevities of conifers in subalpine forests. Oikos 118, 1130–1138. doi: https://doi.org/10.1111/j.1600-0706.2009.17592.x.

Biondi, F., and Qeadan, F. (2008). A Theory-Driven Approach to Tree-Ring Standardization: Defining the Biological Trend from Expected Basal Area Increment. Tree-Ring Res. 64, 81–96. doi: 10.3959/2008-6.1.

Boden, S., Kahle, H.-P., Wilpert, K. von, and Spiecker, H. (2014). Resilience of Norway spruce (Picea abies (L.) Karst) growth to changing climatic conditions in Southwest Germany. For. Ecol. Manage. 315, 12–21. doi: https://doi.org/10.1016/j.foreco.2013.12.015.

Cailleret, M., Jansen, S., Robert, E. M. R., Desoto, L., Aakala, T., Antos, J. A., et al. (2017). A synthesis of radial growth patterns preceding tree mortality. Glob. Chang. Biol. 23, 1675–1690. doi: 10.1111/gcb.13535.

Cailleret, M., Dakos, V., Jansen, S., Robert, E. M. R., Aakala, T., Amoroso, M. M., et al. (2019). Early-Warning Signals of Individual Tree Mortality Based on Annual Radial Growth. Front. Plant Sci. 9, 1964. Available at: https://www.frontiersin.org/article/10.3389/fpls.2018.01964.

Camarero, J. J., Gazol, A., Sangueesa-Barreda, G., Oliva, J., and Vicente-Serrano, S. M. (2015). To die or not to die: early warnings of tree dieback in response to a severe drought. J. Ecol. 103, 44–57. doi: 10.1111/1365-2745.12295.

Clark, J. S., Bell, D. M., Kwit, M., Stine, A., Vierra, B., and Zhu, K. (2012). Individual-scale inference to anticipate climate-change vulnerability of biodiversity. Philos. Trans. R. Soc. B-Biological Sci. 367, 236–246.

Díaz-Martínez, P., Ruiz-Benito, P., Madrigal-González, J., Gazol, A., and Andivia, E. (2023). Positive effects of warming do not compensate growth reduction due to increased aridity in Mediterranean mixed forests. Ecosphere 14, e4380. doi: https://doi.org/10.1002/ecs2.4380.

Dobbertin, M. (2005). Tree growth as indicator of tree vitality and of tree reaction to environmental stress: a review. Eur. J. For. Res. 124, 319–333.

Ferrenberg, S., Kane, J. M., and Mitton, J. B. (2014). Resin duct characteristics associated with tree resistance to bark beetles across lodgepole and limber pines. Oecologia 174, 1283–1292. doi: 10.1007/s00442-013-2841-2.

Fritts, H.C. (1976). Tree rings and climate. Academic Press, New York.

Galiano, L., Martinez-Vilalta, J., and Lloret, F. (2010). Drought-Induced Multifactor Decline of Scots Pine in the Pyrenees and Potential Vegetation Change by the Expansion of Co-occurring Oak Species. Ecosystems 13, 978–991. doi: 10.1007/s10021-010-9368-8

Gazol, A., and Camarero, J. J. (2022). Compound climate events increase tree drought mortality across European forests. Sci. Total Environ. 816, 151604. doi: https://doi.org/10.1016/j.scitotenv.2021.151604.

Gea-Izquierdo, G., Viguera, B., Cabrera, M., and Cañellas, I. (2014). Drought induced decline could portend widespread pine mortality at the xeric ecotone in managed mediterranean pine-oak woodlands. For. Ecol. Manage. 320, 70–82. doi: https://doi.org/10.1016/j.foreco.2014.02.025.

Gea-Izquierdo, G., Férriz, M., García-Garrido, S., Aguín, O., Elvira-Recuenco, M., Hernandez-Escribano, L., et al. (2019). Synergistic abiotic and biotic stressors explain widespread decline of Pinus pinaster in a mixed forest. Sci. Total Environ. 685, 963–975. doi: https://doi.org/10.1016/j.scitotenv.2019.05.378.

Greenwood, S., Ruiz-Benito, P., Martínez-Vilalta, J., Lloret, F., Kitzberger, T., Allen, C. D., et al. (2017). Tree mortality across biomes is promoted by drought intensity, lower wood density and higher specific leaf area. Ecol. Lett. 20, 539–553. doi: https://doi.org/10.1111/ele.12748.

Hammond, W. M., Williams, A. P., Abatzoglou, J. T., Adams, H. D., Klein, T., López, R., et al. (2022). Global field observations of tree die-off reveal hotter-drought fingerprint for Earth’s forests. Nat. Commun. 13, 1761. doi: 10.1038/s41467-022-29289-2.

Hampe, A., and Petit, R. J. (2005). Conserving biodiversity under climate change: the rear edge matters. Ecol. Lett. 8, 461–467. doi: 10.1111/j.1461-0248.2005.00739.x

Hampe, A., and Jump, A. S. (2011). “Climate Relicts: Past, Present, Future,” in Annual Review of Ecology, Evolution, and Systematics, Vol 42, 313–333. doi: 10.1146/annurev-ecolsys-102710-145015

Hartmann, H., Moura, C. F., Anderegg, W. R. L., Ruehr, N. K., Salmon, Y., Allen, C. D., et al. (2018). Research frontiers for improving our understanding of drought-induced tree and forest mortality. New Phytol. 218, 15–28. doi: https://doi.org/10.1111/nph.15048.

Hereş, A.M., Martínez-Vilalta, J., and Claramunt López, B. (2012). Growth patterns in relation to drought-induced mortality at two Scots pine (Pinus sylvestris L.) sites in NE Iberian Peninsula. Trees 26, 621–630. doi: 10.1007/s00468-011-0628-9.

Herguido, E., Granda, E., Benavides, R., García-Cervigón, A. I., Camarero, J. J., and Valladares, F. (2016). Contrasting growth and mortality responses to climate warming of two pine species in a continental Mediterranean ecosystem. For. Ecol. Manage. 363, 149–158. doi: https://doi.org/10.1016/j.foreco.2015.12.038.

Herrero, A., Castro, J., Zamora, R., Delgado-Huertas, A., and Querejeta, J. I. (2013). Growth and stable isotope signals associated with drought-related mortality in saplings of two coexisting pine species. Oecologia 173, 1613–1624. doi: 10.1007/s00442-013-2707-7.

Herrero, A., and Zamora, R. (2014). Plant responses to extreme climatic events: A field test of resilience capacity at the southern range edge. PLoS One 9, e87842. doi: 10.1371/journal.pone.0087842.

Hughes, R. F., Archer, S. R., Asner, G. P., Wessman, C. A., McMurtry, C., Nelson, J. I. M., et al. (2006). Changes in aboveground primary production and carbon and nitrogen pools accompanying woody plant encroachment in a temperate savanna. Glob. Chang. Biol. 12, 1733–1747. doi: https://doi.org/10.1111/j.1365-2486.2006.01210.x.

IPCC (2021) Climate Change 2021: The Physical Science Basis. Contribution of Working Group I to the Sixth Assessment Report of the Intergovernmental Panel on Climate Change [Masson-Delmotte, V., Zhai, P., Pirani, A., Connors, S.L., Péan, C., Berger, S., et al. (eds.)]. Cambridge University Press, Cambridge, United Kingdom and New York, NY, USA, 2391 pp. doi:10.1017/9781009157896.

Linares, J. C., Camarero, J. J., and Carreira, J. A. (2010). Competition modulates the adaptation capacity of forests to climatic stress: insights from recent growth decline and death in relict stands of the Mediterranean fir Abies pinsapo. J. Ecol. 98, 592–603. doi: 10.1111/j.1365-2745.2010.01645.x

Lorimer, C. G. (1983). Tests of age-independent competition indices for individual trees in natural hardwood stands. For. Ecol. Manage. 6, 343–360. doi: https://doi.org/10.1016/0378-1127(83)90042-7.

Madrigal-González, J., Andivia, E., Zavala, M. A., Stoffel, M., Calatayud, J., Sánchez-Salguero, R., et al. (2018). Disentangling the relative role of climate change on tree growth in an extreme Mediterranean environment. Sci. Total Environ. 642, 619–628. doi: https://doi.org/10.1016/j.scitotenv.2018.06.064.

Marqués, L., Camarero, J. J., Zavala, M. A., Stoffel, M., Ballesteros-Cánovas, J. A., Sancho-García, C., et al. (2021). Evaluating tree-to-tree competition during stand development in a relict Scots pine forest: how much does climate matter? Trees 35, 1207–1219. doi: 10.1007/s00468-021-02109-8.

Martín-Moreno, C., Fidalgo Hijano, C., Martín Duque, J. F., González Martín, J. A., Zapico Alonso, I., and Laronne, J. B. (2014). The Ribagorda sand gully (east-central Spain): Sediment yield and human-induced origin. Geomorphology 224, 122–138. doi: https://doi.org/10.1016/j.geomorph.2014.07.013.

Martínez-Vilalta, J., and Pinol, J. (2002). Drought-induced mortality and hydraulic architecture in pine populations of the NE Iberian Peninsula. For. Ecol. Manage. 161, 247–256.

McDowell, N. G., Beerling, D. J., Breshears, D. D., Fisher, R. A., Raffa, K. F., and Stitt, M. (2011). The interdependence of mechanisms underlying climate-driven vegetation mortality. Trends Ecol. Evol. 26, 523–532. doi:10.1016/j.tree.2011.06.003

McDowell, N. G., Sapes, G., Pivovaroff, A., Adams, H. D., Allen, C. D., Anderegg, W. R. L., et al. (2022). Mechanisms of woody-plant mortality under rising drought, CO2 and vapour pressure deficit. Nat. Rev. Earth Environ. 3, 294–308. doi: 10.1038/s43017-022-00272-1.

Millar, C. I., and Stephenson, N. L. (2015). Temperate forest health in an era of emerging megadisturbance. Science 349, 823–826. doi: 10.1126/science.aaa9933.

Muñoz-Gálvez, F. J., Herrero, A., Esther Pérez-Corona, M., and Andivia, E. (2021). Are pine-oak mixed stands in Mediterranean mountains more resilient to drought than their monospecific counterparts? For. Ecol. Manage. 484. doi: 10.1016/j.foreco.2021.118955.

Ogle, K., Whitham, T. G., and Cobb, N. S. (2000). Tree-ring variation in pinyon predicts likelihood of death following severe drought. Ecology 81, 3237–3243.

Pedersen, B. S. (1998). Modeling tree mortality in response to short- and long-term environmental stresses. Ecol. Modell. 105, 347–351.

Peñuelas, J., Ogaya, R., Boada, M., and S. Jump, A. (2007). Migration, invasion and decline: changes in recruitment and forest structure in a warming-linked shift of European beech forest in Catalonia (NE Spain). Ecography 30, 829–837. doi: https://doi.org/10.1111/j.2007.0906-7590.05247.x.

Pinheiro, J., Bates, D., DebRoy, S., and Sarkar, D. R. (2018). Linear and Nonlinear Mixed Effects Models. R package version 3.1-131.1 http://CRAN.R-project.org/package=nlme

R Core Team (2022). R: A language and environment for statistical computing. R Foundation for Statistical Computing, Vienna, Austria. URL: https://www.R-project.org/.

Rowland, L., da Costa, A. C. L., Galbraith, D. R., Oliveira, R. S., Binks, O. J., Oliveira, A. A. R., et al. (2015) Death from drought in tropical forests is triggered by hydraulics not carbon starvation. Nature 528, 119–122. doi: 10.1038/nature15539.

Royer, P. D., Cobb, N. S., Clifford, M. J., Huang, C.-Y., Breshears, D. D., Adams, H. D., et al. (2011). Extreme climatic event-triggered overstorey vegetation loss increases understorey solar input regionally: primary and secondary ecological implications. J. Ecol. 99, 714–723. doi: https://doi.org/10.1111/j.1365-2745.2011.01804.x.

Ruiz-Benito, P., Lines, E. R., Gómez-Aparicio, L., Zavala, M. A., and Coomes, D. A. (2013). Patterns and Drivers of Tree Mortality in Iberian Forests: Climatic Effects Are Modified by Competition. PLoS One 8, e56843. https://doi.org/10.1371/journal.pone.0056843.

Ruiz-Benito, P., Ratcliffe, S., Zavala, M. A., Martínez-Vilalta, J., Vilà-Cabrera, A., Lloret, F., et al. (2017). Climate- and successional-related changes in functional composition of European forests are strongly driven by tree mortality. Glob. Chang. Biol. 23, 4162–4176. doi: https://doi.org/10.1111/gcb.13728.

Sangüesa-Barreda, G., Linares, J. C., and Camarero, J. J. (2015). Reduced growth sensitivity to climate in bark-beetle infested Aleppo pines: Connecting climatic and biotic drivers of forest dieback. For. Ecol. Manage. 357, 126–137. doi: https://doi.org/10.1016/j.foreco.2015.08.017.

Schneider, C. A., Rasband, W. S., and Eliceiri, K. W. (2012). NIH Image to ImageJ: 25 years of image analysis. Nat. Methods 9, 671–675. doi: 10.1038/nmeth.2089.

Seidl, R., Fernandes, P. M., Fonseca, T. F., Gillet, F., Jönsson, A. M., Merganičová, K., et al. (2011). Modelling natural disturbances in forest ecosystems: a review. Ecol. Modell. 222, 903–924. doi: https://doi.org/10.1016/j.ecolmodel.2010.09.040.

Shestakova, T. A., Gutiérrez, E., Kirdyanov, A. V, Camarero, J. J., Génova, M., Knorre, A. A., et al. (2016). Forests synchronize their growth in contrasting Eurasian regions in response to climate warming. Proc. Natl. Acad. Sci. 113, 662–667. doi: 10.1073/pnas.1514717113

Steckel, M., del Río, M., Heym, M., Aldea, J., Bielak, K., Brazaitis, G., et al. (2020). Species mixing reduces drought susceptibility of Scots pine (Pinus sylvestris L.) and oak (Quercus robur L., Quercus petraea (Matt.) Liebl.) – Site water supply and fertility modify the mixing effect. For. Ecol. Manage. 461, 117908. doi: https://doi.org/10.1016/j.foreco.2020.117908.

Sumida, A., Miyaura, T., and Torii, H. (2013). Relationships of tree height and diameter at breast height revisited: analyses of stem growth using 20-year data of an even-aged Chamaecyparis obtusa stand. Tree Physiol. 33, 106–118. doi: 10.1093/treephys/tps127.

Tejedor, E., Serrano-Notivoli, R., de Luis, M., Saz, M. A., Hartl, C., St. George, S., et al. (2020). A global perspective on the climate-driven growth synchrony of neighbouring trees. Glob. Ecol. Biogeogr. 29, 1114–1125. doi: https://doi.org/10.1111/geb.13090.

Thornthwaite, C.W. (1948) An approach toward a rational classification of climate. Geogr. Rev. 38:55–94

Vila-Cabrera, A., Martinez-Vilalta, J., Galiano, L., and Retana, J. (2013). Patterns of Forest Decline and Regeneration Across Scots Pine Populations. Ecosystems 16, 323–335. doi: 10.1007/s10021-012-9615-2

Williams, A.P., Allen, C. D., Macalady, A. K., Griffin, D., Woodhouse, C. A., Meko, D. M., et al. (2013). Temperature as a potent driver of regional forest drought stress and tree mortality. Nat. Clim. Chang. 3, 292–297. doi: 10.1038/nclimate1693.

Xiong, Y., D’Atri, J. J., Fu, S., Xia, H., and Seastedt, T. R. (2011). Rapid soil organic matter loss from forest dieback in a subalpine coniferous ecosystem. Soil Biol. Biochem. 43, 2450–2456. doi: https://doi.org/10.1016/j.soilbio.2011.08.013.

Yamaguchi, D. K. (1991). A simple method for cross-dating increment cores from living trees. Can. J. For. Res. 21, 414–416. doi: 10.1139/x91-053.

Zuur, A. F., Ieno, E. N., Walker, N., Saveliev, A. A., Smith, G. M. (2009) Mixed effects models and extensions in ecology with R. Springer, United Kingdom. 579 pp.

